# Let’s Do the Time Warp Again: Non-linear time series matching as a tool for sequentially structured data in ecology

**DOI:** 10.1101/2021.04.19.440490

**Authors:** Jens C. Hegg, Brian P. Kennedy

## Abstract

Ecological patterns are often fundamentally chronological. However, generalization of data is necessarily accompanied by a loss of detail or resolution. Temporal data in particular contains information not only in data values but in the temporal structure, which is lost when these values are aggregated to provide point estimates. Dynamic Time Warping (DTW) is a time series comparison method that is capable of efficiently comparing series despite temporal offsets that confound other methods. The DTW method is both efficient and remarkably flexible, capable of efficiently matching not only time series but any sequentially structured dataset, which has made it a popular technique in machine learning, artificial intelligence, and big data analytical tasks. DTW is rarely used in ecology despite the ubiquity of temporally structured data. As technological advances have increased the richness of small-scale ecological data, DTW may be an attractive analysis technique because it is able to utilize the additional information contained in the temporal structure of many ecological datasets. In this study we use an example dataset of high-resolution fish movement records obtained from otolith microchemistry to compare traditional analysis techniques with DTW clustering. Our results suggest that DTW is capable of detecting subtle behavioral patterns within otolith datasets which traditional data aggregation techniques cannot. These results provide evidence that the DTW method may be useful across many of the temporal data types commonly collected in ecology, as well other sequentially ordered “pseudo time series” data such as classification of species by shape.

**Keywords:** classification, cluster analysis, data generalization, DTW, dynamic time warping, otolith chemistry, time series

## Introduction

The study of ecology is fundamentally chronological, and the challenges ecologists face with the collection and analysis of their data often reflects this temporal nature (Wolkovich et al. 2014). Populations rise and fall over years. Climate, as well as the rates of predation, parasitism, and competition vary across time, affecting behavior, survival, and reproduction of populations. Analyzing and modeling this data often requires translating data collected at small scales into meaningful metrics that can explain larger phenomenon. This translation, however, inevitably results in loss of information though loss of detail and specificity (Levin 1992).

Levin (1992) argues convincingly that simplifying data from the individual to the ecosystem scale should be done with the goal of thoughtfully preserving “minimal sufficient detail” to inform models at larger scales. However, as technology drives increases in the volume and richness of data at the individual and local scale (Hampton et al. 2009, Laurance et al. 2016), it is reasonable to assume that small-scale data may now contain more meaningful data. Analyses that better summarize information-rich data could increase the meaning of aggregated data. Recent advances in time series analysis techniques may be an example of just such a technique (Aghabozorgi et al. 2015). Time series data contains information not just in the values of the data, but in the order of those values (Chatfield 2003, Cressie and Wikle 2011). Many analysis techniques collapse these data into discrete time-points, or overall descriptive statistics, in the process removing temporal structure that may be more useful than we realize. New time series analysis tools that have gained prominence in other fields may allow more nuanced treatment of time series data in ecology, decreasing the loss of information in time series data due to aggregation.

One of the most popular time series techniques is Dynamic Time Warping (DTW). DTW distance is a distance measure (similar to the familiar Euclidian or Mahalanobis distance measures) which describes the similarity of two time series, or any dataset which can be expressed sequentially. It was first developed as a method to match sounds in speech recognition, where the speed and accent of speakers can vary despite the word or phrase being the same, creating phase shifts that are difficult to match using most distance metrics (Sakoe and Chiba 1978, Myers and Rabiner 1981). DTW excels at matching similar time series which vary temporally and has been shown to be fast, and highly efficient for classification (Al-Naymat et al. 2009, Rakthanmanon et al. 2012). The technique is flexible and can be applied using existing clustering and classification methods (Mueen and Keogh 2016, Sarda-Espinosa 2017).

The DTW distance describes the Euclidean distance between two time series, after first “warping” them into alignment. An accessible and concise explanation of the mathematical basis and statistical applications of the technique is available in Ratanamahatana and Keogh (2004). Briefly, DTW finds the optimal path across the matrix created by matching each point in a time series with each point of a comparison series (Figure 1A). This path is found using dynamic programming to minimize a cost function for each sequential step across the matrix, essentially finding the path which matches the most similar points in each series. In the case of two identical time series, the least cost path would be a perfect diagonal. For misaligned series, the algorithm matches each point in the two time series with a one-to-many approach, warping the temporal dimension to match the two series (Figure 1B). This warping technique allows time series to be compared after correction for temporal differences that would otherwise skew a Euclidean distance measure.

**Figure 1-.**
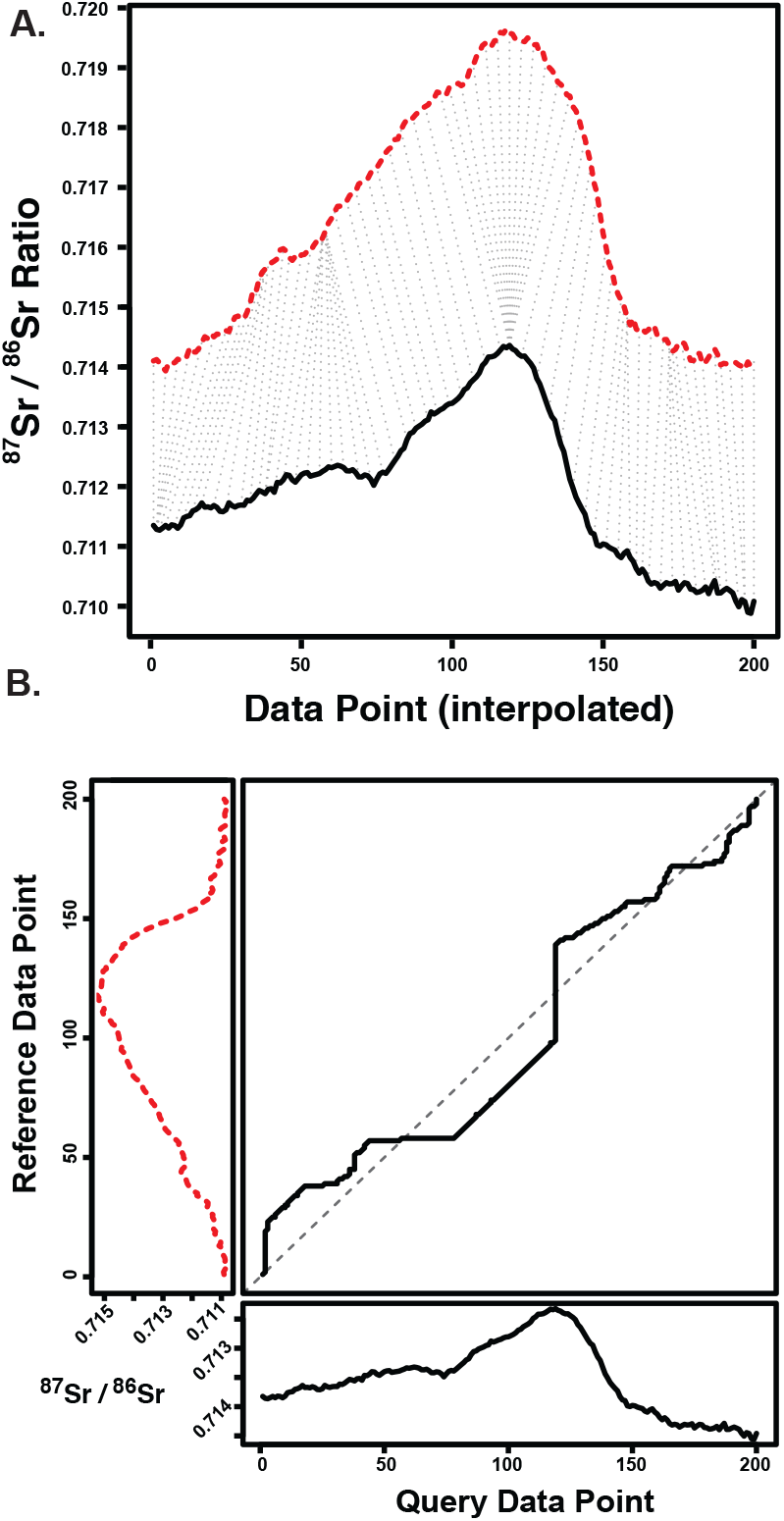
Two otolith ^87^Sr/^86^Sr transects of two fish caught in the LGR reach of the Snake River are shown. Dynamic Time Warping computes the amount of warping on the temporal (x) axis needed to optimally align two series (A). Dotted grey lines show matching points along these series computed by DTW (a subsample of matching points is shown and transects are offset by 0.004 to improve clarity). The optimal warping path (B) is shown between the two time series. Transects are re-interpolated to 200 cells but were not z-normalized prior to comparison.

Despite the increasing popularity of DTW in big-data mining (Keogh and Pazzani 2000, Sakurai et al. 2015), artificial intelligence & robotics (Xu et al. 2014, Cheng et al. 2015), economics (Lee et al. 2012, Wang et al. 2012), healthcare (Ortiz et al. 2016), and speech recognition (Pi-Yun Chen et al. 2015) these techniques have only been applied in a few cases to ecological data (Debeljak et al. 2010, Cope and Remagnino 2012, Stathopoulos et al. 2014, Tan et al. 2015, Jouary et al. 2016, Baumann et al. 2017, Weideman et al. 2017).

DTW may be useful for many types of ecological data because much of the data that ecologists collect has a temporal component; for example, spatial location data of tagged animals, mark-recapture data, trends in population density, and the timing of spring leaf-out, all are either inherently temporal or could be thought of as a time series. In fact, the use of DTW to analyze “pseudo time series,” sequentially ordered data which is not temporal but can be thought of as such for the sake of analysis, has potentially wide application in ecology. Identification of plant or fish species based on shape, or animal movement patterns, are three examples in the literature (Ueno et al. 2006, Cope and Remagnino 2012, Jouary et al. 2016). Even genetics data can be coerced into a time series format for use with time series methods (Rakthanmanon et al. 2012). In many of these cases it is useful to determine the similarity or difference between the structure of data; for example, to classify streams by the characteristics of their hydrograph, or to track the timing of phenological events across decades. Time series clustering tools like DTW provide methods which can efficiently cluster similar time series using all the information contained in the chronological structure of the data, avoiding some of the problems associated with data aggregation.

One example of temporally structured data for which DTW may provide analytical advantages is the life-time chemical records obtained from fish otoliths (or ear stones). Over the same period that time series clustering methods have matured, the microchemical analysis of fish otoliths has taken similarly large strides as an ecological tool, becoming an example of a dramatic increase in data richness at the individual scale (Campana 2005, Secor 2010, Walther 2019). With calcium carbonate rings laid down daily, the otolith is a natural temporal record of the environment and life-history of a fish (Campana and Neilson 1985). Otoliths record ambient chemistry which can be used to reconstruct an individual’s movements, life-history strategies, and environmental conditions through the life of a fish with remarkable precision (Kennedy et al. 1997, 2002, Campana and Thorrold 2001, Hamann and Kennedy 2012, Limburg et al. 2013). Studies now often including multiple chemical and isotopic tracers, each able to reconstruct different multiple facets of a fish’s life-history (Walther and Limburg 2012, Hegg et al. 2018). These new techniques create information-rich, time series data of a fish from birth to death (Figure 2A).

**Figure 2-.**
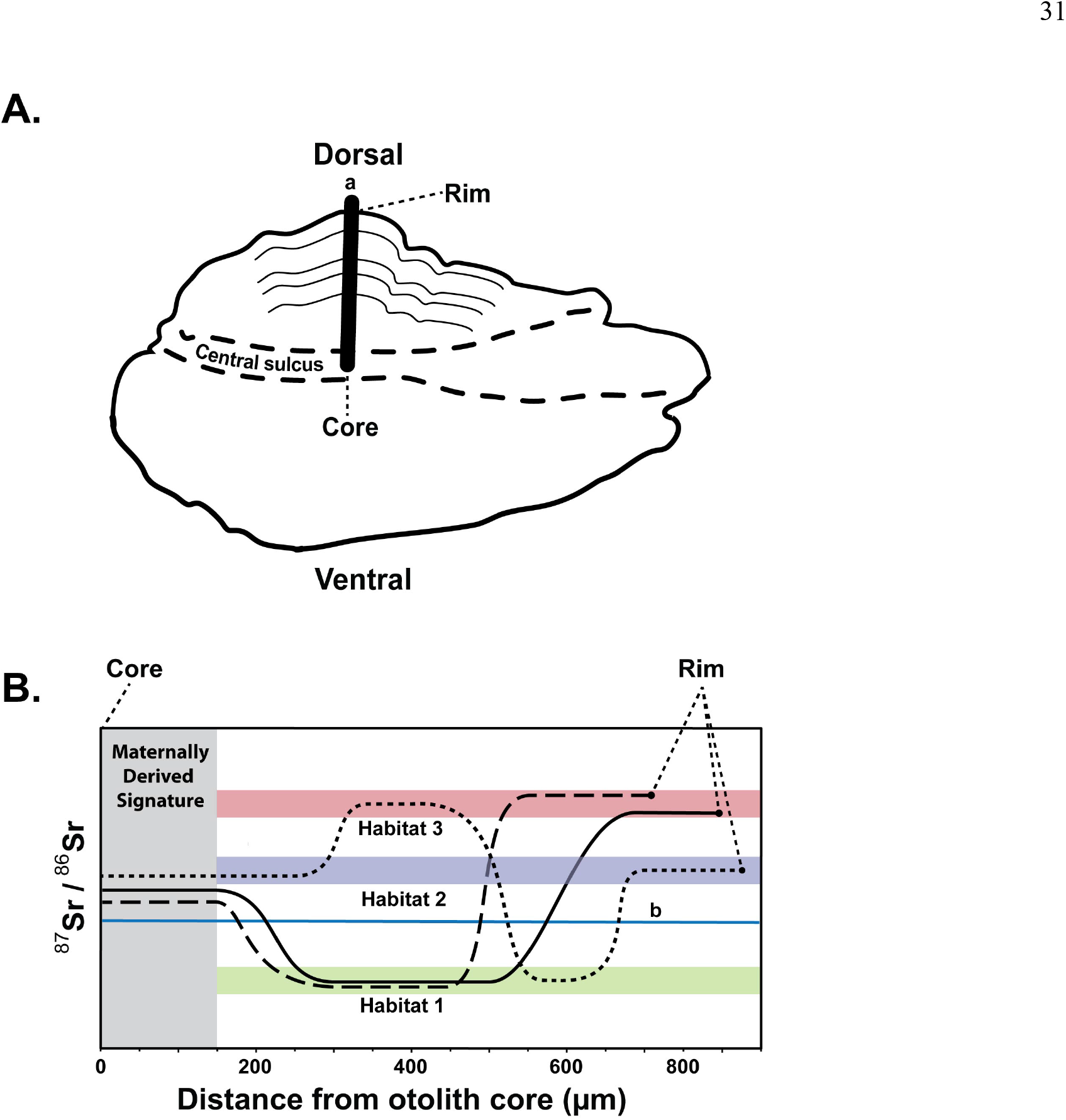
Otoliths (A) are laser ablated on a path 90° from the sulcus (a). This analysis creates a temporally structured dataset for each fish, representing the chemistry of the rivers they inhabited throughout their life. These otolith chemical transects (B) are represented with microns from the core of the otolith on the temporal (x) axis. Movement timing, growth and location combine to form the shape of the ^87^Sr/^86^Sr curve as a fish moves through different habitats. Otolith ^87^Sr/^86^Sr transects for to hypothetical fish inhabiting the same two habitats, habitats 1 and 3, are shown (solid and heavy dashed black lines). The otolith transects for these fish which experienced the same habitats are phase-shifted on the temporal axis, a condition which is not controlled in Euclidean distance time series matching. the flexible temporal dimension in the DTW method allows for matching these transects while distinguishing other life-history shapes (fine dashed black line). The global marine ^87^Sr/^86^Sr signature is shown (b) for reference.

While the resolution of data extracted from otoliths has increased dramatically, analysis techniques have not taken advantage of the increased information density of high-resolution time series datasets. In most cases data from periods of interest in the fish’s life is aggregated, creating a chemical index which can be analyzed as a discrete value, or a vector of values in the multivariate case (Barnett-Johnson et al. 2010, Hegg et al. 2013a, Garcez et al. 2014, Hegg et al. 2018). While this is a valid approach, it risks ignoring or obscuring valuable information contained in the time series structure itself whose shape is affected by growth, movement, and ontogeny.

In this paper, we provide an example of how DTW can be used to determine natal origin and life-history from a large dataset of known-origin juvenile Chinook salmon otolith transect data. We demonstrate the ability of DTW to cluster fish using univariate and multivariate ^87^Sr/^86^Sr data. Next, we compare these results to a more conventional, model-based discriminate function analysis using aggregated multivariate data, and we demonstrate how the two techniques could be paired to potentially improve location discrimination. We then present the use of nearest-neighbor classification to extend DTW clustering to classify unknown fish. Finally, we discuss the utility of DTW methods in ecology more broadly given our results.

## Methods

### Study Species

Snake River fall Chinook salmon are a threatened population of fall-spawning *Oncorhynchus tshawytscha* in the Snake River of Idaho, a major tributary to the Columbia River in the northwestern United States (Figure 3). The population is notable for a recent shift in juvenile life-history strategy which recent research suggests is hereditary and an example of contemporary life-history evolution (Williams et al. 2008, Waples et al. 2017). These changes are thought to be driven by anthropogenic changes related to ten hydropower dams which have blocked the majority of historical spawning habitat and created significant changes in the hydrograph and river conditions throughout their current range (Connor et al. 2005, Hegg et al. 2013a, Connor et al. 2016).

**Figure 3-.**
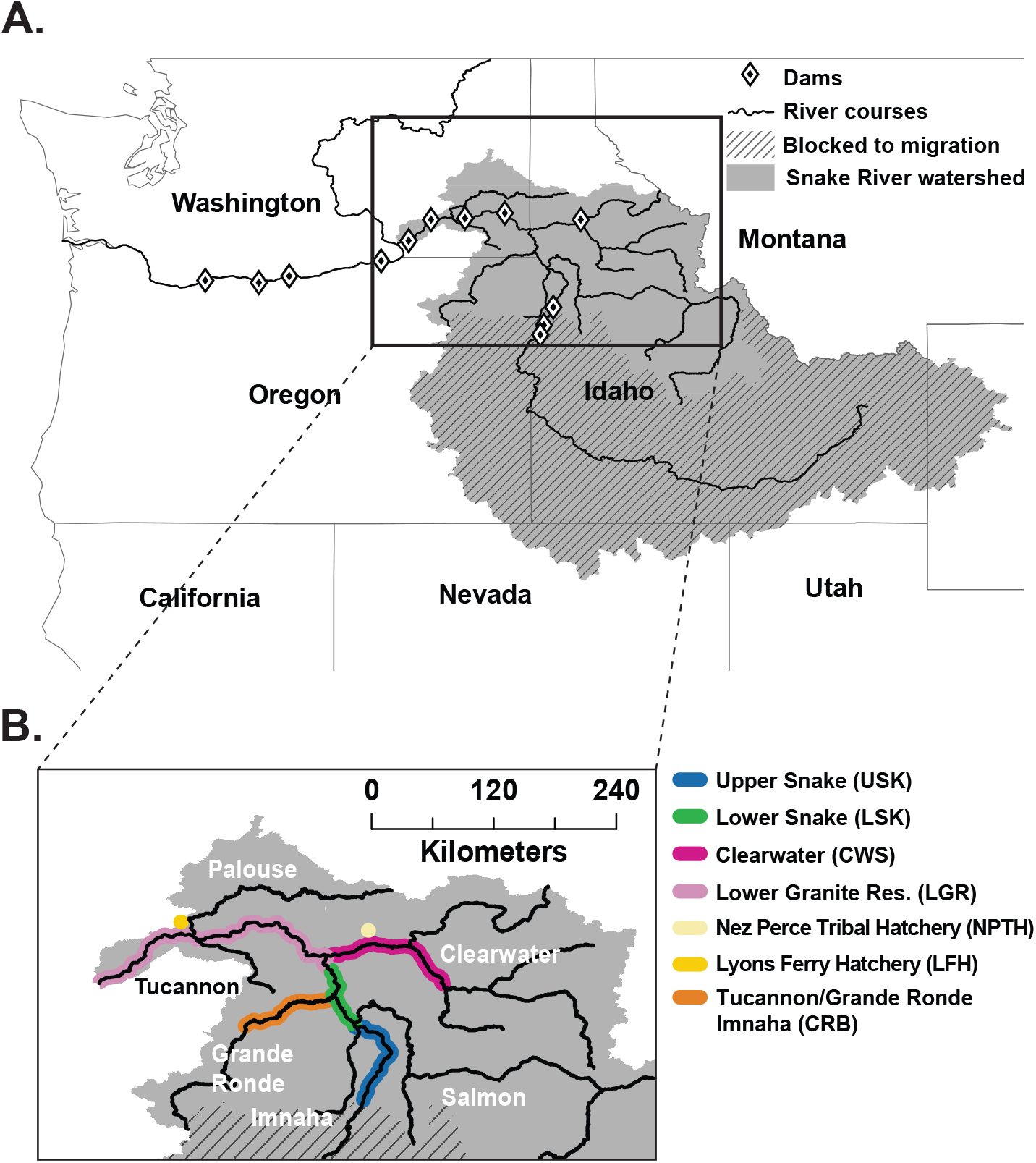
Snake River fall Chinook salmon inhabit the Snake River (a) in the US States of Idaho, Washington and Oregon. The extent of spawning for known origin fish in our study (b) is highlighted. The location of the two hatcheries in the basin are noted with colored dots. The Tucannon, Grande Ronde, and Salmon Rivers were not sampled for juveniles and produce a very small percentage of the wild fish in the basin.

The geology of the major spawning tributaries in this watershed is diverse both in age and rock type, creating significant differences in water chemistry between the major spawning areas (Hegg et al. 2013b). Ongoing water sampling throughout the basin has shown that ^87^Sr/^86^Sr signatures in the four main spawning areas of the basin are distinct, and that discriminate function analysis using ^87^Sr/^86^Sr can be used to determine the locations of juvenile and adult fish using otolith chemistry (Hegg et al. 2013a, 2018).

## Otolith Collection and Analysis

The data used in this study consists of otoliths collected from known-origin, juvenile fall Chinook salmon from throughout their range in the Snake River basin as part of a prior otolith study (Hegg et al. 2018). Juveniles were collected at three locations from 2009 to 2014 (n=376) as part of population surveys conducted across the spawning areas in the Snake, Grande Ronde and Clearwater Rivers by United States Fish and Wildlife Service, Nez Perce Tribe Department of Fisheries Resource Management, and USGS. Samples were also obtained from the two hatcheries producing Fall Chinook in the basin, Lyons Ferry Hatchery and the Nez Perce Tribal Hatchery. Some fish were tagged with passive integrated transponder (PIT) tags and released, then recaptured weeks later when their tag was detected at Lower Granite Dam, the first dam fish encounter on their path downstream.

Otolith samples were collected and processed using established procedures for otolith analysis (Secor et al. 1992, Hegg et al. 2013a). Detailed methods for this dataset, including laser ablation and ICP-MS information, are available in Hegg et al. (2018). Continuous chemical transects from the core (birth) to the edge (death) were collected from each otolith, creating a sequential chemical record throughout the life of the fish (Figure 2A and B). This was done using using a New Wave UP-213 laser ablation sampling system. This system was coupled with a Thermo Scientific Neptune multi-collector inductively coupled plasma mass-spectrometer (ICP-MS) for ^87^Sr/^86^Sr ratio. The ablation system was coupled with a Thermo Scientific Element2 ICP-MS to measure elemental concentrations of calcium (^43^Ca), strontium (^86^Sr), barium (^138^Ba), Magnesium (^25^Mg), and Manganese (^55^Mn). Elemental measurements were calculated as ratios to calcium and expressed as mm/mol following Hegg et al. (2018).

### Multivariate Discriminate Function Classification

A model-based discriminant function was created to classify fish to known location. The goal was to develop a robust classification which could than later be applied to unknown adult fish to inform ecology and management of the population.

Within our dataset, juveniles were assigned to a known origin based on the location of capture. Juveniles which were captured and sacrificed during beach seine sampling were assigned to the river reach in which they were captured; the Upper Snake River (USK), the Lower Snake River (USK), the Clearwater River (CWS), or the Grande Ronde River (GR). Fish that were PIT tagged and recaptured at Lower Granite Dam were assigned a second known location in Lower Granite Reservoir (LGR). Therefore, it was possible for a fish to have both a known natal location and a known downstream location. Some fish were caught in the dam forebay as a part of prior studies and their natal location was not known, although their migration timing suggested Clearwater River origin. These fish were assigned only a known downstream location of LGR. Juveniles obtained from Lyons Ferry Hatchery (LFH) and Nez Perce Tribal Hatchery (NPTH) were assigned to these natal locations respectively.

We defined the natal signature as the average of the chemistry between 300μm and 400μm from the core of the otolith, creating a five-element vector of chemical signatures for each fish. This was based on prior research showing this to be the beginning of stable juvenile signatures (Hegg et al. 2018). Fish arriving at Lower Granite Dam can be moving downstream quickly and may have only recently equilibrated. Therefore, we averaged the signatures from only the outer 50μm from the edge to obtain the downstream signature. Fish captured at LGR with a ^87^Sr/^86^Sr signature far removed from that of LGR were assumed to be fast moving migrants, fish moving too fast to have equilibrated to the surrounding water. These were removed to maintain a consistent training set for the discriminant function.

Classifying fish to location was done using a model-based discriminant function using ^87^Sr/^86^Sr, Sr/Ca, Ba/Ca, Mn/Ca, and Mg/Ca as independent variables and known location as the classifier. We used the {mclust} package (version 5.2) for R to build a model based discriminant function (Fraley and Raftery 2007, Scrucca et al. 2016). The dataset was randomly split into a training set (80%) and test set (20%), with the training-set used to construct the discriminate function. Related river reaches were combined successively until an acceptable misclassification rate was achieved. The final discriminate function was then applied to the test-set to quantify its performance on unknown data.

### Dynamic Time Warping Cluster Analysis

Time series clustering using DTW distance was used to identify groups of similar fish based on the shape of their otolith transects. The DTW algorithm is extremely sensitive to variation in mean, requiring all time series to be normalized (Keogh and Kasetty 2002, Rakthanmanon et al. 2012). In the case of fish life-history transects this normalization can be problematic, as the absolute mean of the ^87^Sr/^86^Sr ratio is meaningful as a marker of fish location, and once normalized to a mean of zero and unit variance it is possible for the shape of two otolith transects from different rivers to look alike, and thus cluster together despite being meaningfully different.

To remove the location ambiguity created by z-normalization we used a two-step clustering process to partition the transects by mean, and then sub-cluster by DTW distance. The mean of each ^87^Sr/^86^Sr transect was calculated, means data was scaled, then clustered using model-based clustering in the Mclust package for R. Initial clustering was performed with the intention of finding the minimum number of clusters which cleanly separated the known groups. The clusters obtained from Mclust where then sub-clustered using DTW distance on both univariate ^87^Sr/^86^Sr and multivariate data including the elemental ratios used in the discriminate function classification above.

Prior to DTW clustering, a centered, 60-point, rolling average was used to smooth ^87^Sr/^86^Sr transects. A 10-point rolling average was used to smooth elemental data, as the longer integration time during collection of this data results in smoother data. Transects were then re-interpolated to a length of 200 cells, which allows faster calculation through the implementation of lower bounds without a loss of the ability to accurately match time series (Ratanamahatana and Keogh 2004, Al-Naymat et al. 2009). This length was used as a rounded approximation of the mean length of series in the dataset. The mean length of transects was 173 cells, with a maximum length of 422, a minimum of 61 and a standard deviation of 61 cells. A comparison of selected data before and after interpolation is included in Appendix A.

Clustering was performed on the univariate and multivariate transects using the {dtwclust} package (Sarda-Espinosa 2017) using agglomerative hierarchical clustering {hclust} method in R with Wards distance. A 5% Sakoe-Chiba window was employed to decrease processing effort and limit potentially erroneous warping (Sakoe and Chiba 1978, Ratanamahatana and Keogh 2004, Al-Naymat et al. 2009). The Sakoe-Chiba window limits the amount of deviation from the diagonal when determining the warping path between two time series (Figure 1B). Otolith transects were z-normalized prior to analysis.

Clustering was exploratory, with a goal to cut the dendrogram of each group at a location which minimized the number of clusters while maintaining clusters which were easily interpretable based on the known-origin of the fish within each cluster. Window size was varied from 1% to 100% after the optimal number of groups was found to determine if adjusting the Sakoe-Chiba window (Sakoe and Chiba 1978) affected the stability of the results.

The same clustering approach was repeated for univariate data using the Euclidian distance measure. This was done to test whether DTW was a superior distance metric over the more traditional Euclidean distance which does not take into account temporal warping. Euclidian distance was not performed on the multivariate data as there are no packages which implement multivariate Euclidean distance for time series clustering.

### Combining DTW with Discriminate Function Analysis

In one case the discriminant function was unable to separate two groups of fish from known locations, the USK and LSK, allowing us to test the ability of DTW to separate these indistinguishable groups. We applied hierarchical clustering to the training set data from these confounded groups, reserving the test-set data to test the robustness of the grouping using nearest-neighbor classification. For the clustering step we used a 5% Sakoe-Chiba window and Keogh lower bounds. The effect of window size was tested qualitatively by varying the window from 1% to 100% to test the stability of the clustering results. We cut the dendrogram to create three clusters based on the results of the overall DTW clustering. We then used this cluster solution to predict the cluster membership of the test-set otoliths from these confounded groups using 1-nearest-neighbor classification, to test the stability of these cluster results to unknown data. Comparison to known water chemistry from Hegg et al. (2018) was used to evaluate the veracity of this group membership.

## Results

### Multivariate Discriminate Function Analysis

Initial data exploration indicated that the LGR group, as expected, contained a number of juvenile fish whose signatures had not equilibrated and instead reflected signatures of upstream habitats (n=22). Additionally, the LSK group contained one fish caught in the Lower Snake River and later at LGR which exhibited a very high, Clearwater River signature. These fish were removed to provide a robust training set, under the assumption that adult fish would exhibit a clear signature in these locations, having had time to chemically equilibrate.

Additionally, a group of anomalous life-history transects were identified in the CWS group which did not appear to conform to the known signatures which a Clearwater origin fish would experience, nor did they match the expected patterns or signatures seen in NPTH fish. All of these fish were captured late in the year and could potentially be unmarked hatchery juveniles, erroneously identified Spring Chinook from upriver populations, or an unknown source. To avoid biasing our CWS training set these fish were excluded (n=25).

There was significant overlap in the ^87^Sr/^86^Sr signatures between the USK and LSK groups. Many LSK fish appeared, subjectively, to have originated in the USK and moved very early to the LSK downstream. Evidence of this type of early movement has been observed in the population (Ken Tiffan, USGS, unpublished data). However, without evidence to clearly identify these potentially early-moving juveniles they were kept within their known-origin groupings.

Model based clustering resulted in a model with variable, ellipsoidal and diagonal variance structures for each group. The initial classification attempt resulted in large misclassification errors between the LSK and USK groups. The USK and LSK groups were subsequently combined and the classification was run again. This final training set classification resulted in an absolute training error of 3.6% (n=298), with a 10-fold cross-validation error rate of 12.8% (SE = 3.3%). Classification of the test set (n=74) resulted in an overall classification error rate of 12.2%. (Table 1)

**Table 1.** Classification accuracy of discriminate function training set

|  |  | **Observed** |  |  |  |  |  |
| --- | --- | --- | --- | --- | --- | --- | --- |
|  |  | CWS | CRB | LFH | LGR | NPTH | USK/LSK |
| **Predicted** | CWS | 64 | 0 | 0 | 0 | 0 | 0 |
|  | CRB | 0 | 21 | 0 | 0 | 0 | 0 |
|  | LFH | 0 | 0 | 19 | 0 | 2 | 0 |
|  | LGR | 2 | 0 | 0 | 60 | 0 | 9 |
|  | NPTH | 2 | 0 | 2 | 0 | 21 | 5 |
|  | USK | 0 | 0 | 0 | 2 | 1 | 92 |

### Dynamic Time Warping Cluster Analysis

Initial model-based clustering on the mean of each transect resulted in three clearly defined groupings in the dendrogram which largely corresponded to the three major river systems in the basin (Table 2). The first cluster was made up largely of transects from the Clearwater River (n=101) with high mean ^87^Sr/^86^Sr and is referred to as the Clearwater cluster. Seventeen samples in this group were from other locations, the majority of which were of unknown origin. The second cluster, the Snake River cluster, was a mixed cluster made up of fish from the Upper and Lower Snake Rivers, NPTH and LFH hatcheries with intermediate mean ^87^Sr/^86^Sr transects, The third group was made up entirely of fish from the Grande Ronde River with low ^87^Sr/^86^Sr values.

**Table 2.**
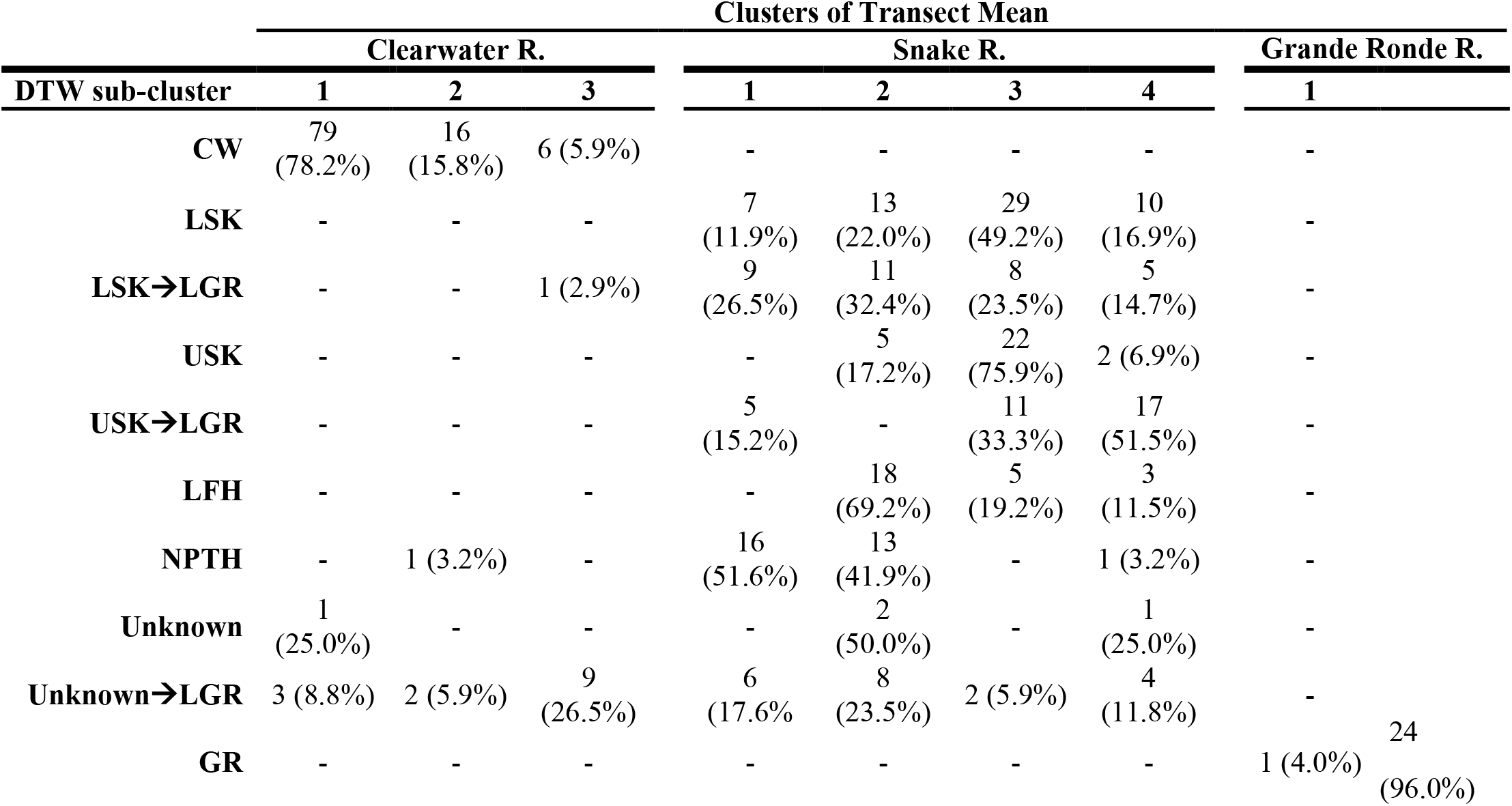
Results of two-level clustering of known-origin juvenile fish using univariate DTW distance. Fish were first clustered by the mean 87Sr/86Sr of the entire transect using k-means, resulting in three broad clusters corresponding to the river of origin (Clearwater River, Snake River, and Grande Ronde River). These clusters were sub-clustered using hierarchical clustering and dynamic time warping distance on the 87Sr/86Sr transect for each otolith. Sample size in each sub-cluster is shown, with the percentage of each known-origin group shown in parentheses.

#### Univariate DTW Sub-clustering

Univariate hierarchical DTW sub-clustering of juvenile ^87^Sr/^86^Sr transects in the Clearwater cluster separated into three clusters (Figure 4A, Table 2). The first cluster was made up of 83 fish with a steeply ascending ^87^Sr/^86^Sr profile consistent with the transition from maternal otolith signature to that of the Clearwater River. The second cluster separated 16 of the fish which were omitted from the discriminate function in the section above due to their anomalous signature. The third cluster consisted of fish with an ^87^Sr/^86^Sr transect that quickly ascended toward the Clearwater and then descended toward the lower signature of the Snake River. The majority of cluster 3 fish were of unknown origin and captured in Lower Granite Reservoir. The remaining fish in cluster 3 exhibited a range of unique patterns.

**Figure 4-.**
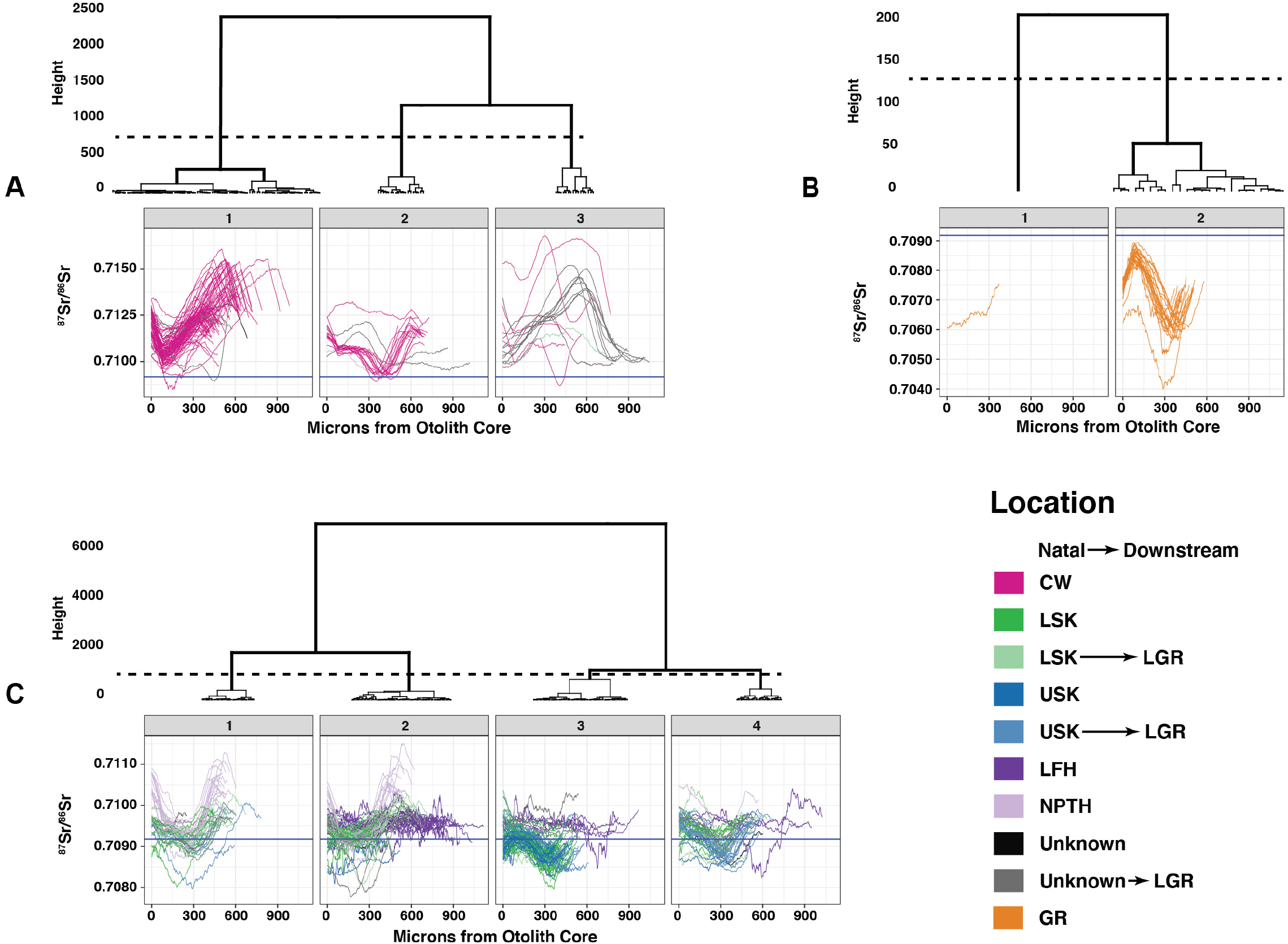
Univariate hierarchical DTW sub-clustering of juvenile fish ^87^Sr/^86^Sr transects are shown. Panels A through C represent the sub-clustering for each cluster based on overall transect mean. Dashed lines show the height the dendrogram was cut to determine the cluster solution. Transects are colored by the known location of the fish. Some fish were captured in their natal location, released, and recaptured downstream in LGR. The blue horizontal line represents the global marine value of ^87^Sr/^86^Sr (0.70918) for reference.

The DTW sub-clustering of the Grande Ronde cluster resulted in two groups, with cluster 1 containing only a single sample with an anomalous, increasing ^87^Sr/^86^Sr transect (Figure 4B, Table 2). The second cluster contained the remainder of the samples, all of which exhibited a steeply declining ^87^Sr/^86^Sr signature followed by a small increase at the end of the transect.

Univariate DTW sub-clustering of the Snake River cluster resulted in four distinct clusters (Figure 4C, Table 2). Subjectively cluster 1 contained fish with a signature beginning near the global marine signature (0.70918) and increasing toward the signature of the USK. This cluster was composed of a majority of fish from NPTH (16) and fish from the LSK which were subsequently captured in LGR (9). Cluster 2 appeared more mixed, with a combination of increasing signatures similar to cluster 1 and a large number of invariant signatures from LFH. The majority of cluster 2 fish were from LFH (18) but large numbers of fish from other locations as well, including LSK (13), NPTH (13), and LSK fish captured in LGR (11). Cluster 3 transects appeared to be dominated by a signature decreasing from the global marine signature toward the USK signature and was dominated by fish from USK (29). This cluster also included a large number of fish captured in the LSK reach (22). Cluster 4 appeared to separate a ^87^Sr/^86^Sr signature decreasing from the global marine signature before rising and crossing the global marine signature to end at the signature of the USK or LGR. The majority of fish in cluster 4 were of USK origin, later captured in LGR (17) with a smaller number originating in the LSK (10).

Overall, the univariate DTW clustering results showed some ability to classify fish, though clusters were not unambiguous. Particularly in the Clearwater River, DTW was able to separate fish which had already been identified as having an anomalous signature (Cluster 2), and to distinguish fish with a likely origin in the Clearwater which were captured further downstream in LGR. Sub-clustering of the Snake River cluster showed some ability to separate hatchery fish (Clusters 1 and 2), and some ability to distinguish patterns in the ^87^Sr/^86^Sr transects which could distinguish fish originating in the USK and LSK despite their later downstream movement which confounded the discriminate function. Varying the size of the Sakoe-Chiba window had very little impact on the DTW results.

Univariate sub-clustering of ^87^Sr/^86^Sr transects using Euclidean distance resulted in very similar clustering results and dendrograms to the DTW results. The most significant difference between the two distance metrics was the ordering of sub-clusters in the Clearwater River. The details of these results are not presented to avoid repetition.

#### Multivariate DTW Sub-clustering

Multivariate, hierarchical DTW sub-clustering resulted in more straightforward sub-clustering results. Multivariate clustering excluded Mg/Ca ratio as a variable. This was done after determining that outliers within the Mg/Ca transects resulted in poor clustering results overall. The interpretability of clustering results improved markedly after removal of Mg/Ca from the dataset.

Within the Clearwater River cluster, DTW identified four distinct cluster groups (Figure 5A, Table 3). Cluster 1 was made up of a majority of fish of unknown natal origin captured in LGR (12) with a single fish from the CW group. Cluster 2 was made up of CW origin fish (20) with a sharply ascending ^87^Sr/^86^Sr transect similar to Clearwater sub-cluster 1in the univariate case. Cluster 3 was made up of 15 samples with the anomalous transect shape which was excluded from the discriminate function and one fish from NPTH. The similarity the NPTH fish classified to the same group supports the idea that these are fish from NPTH which were captured unknowingly. Further, the fish in this cluster were all captured late in 2014 which increases the chances that hatchery juveniles would be included in sampling. The water in which fish are held at NPTH is mixed between well and river sources depending on conditions (Hegg et al. 2018), which could cause differences in the ^87^Sr/^86^Sr curves between cluster 4 and the known-origin NPTH fish in cluster 2. Alternatively, the possibility of an unknown source cannot be eliminated.

**Figure 5-.**
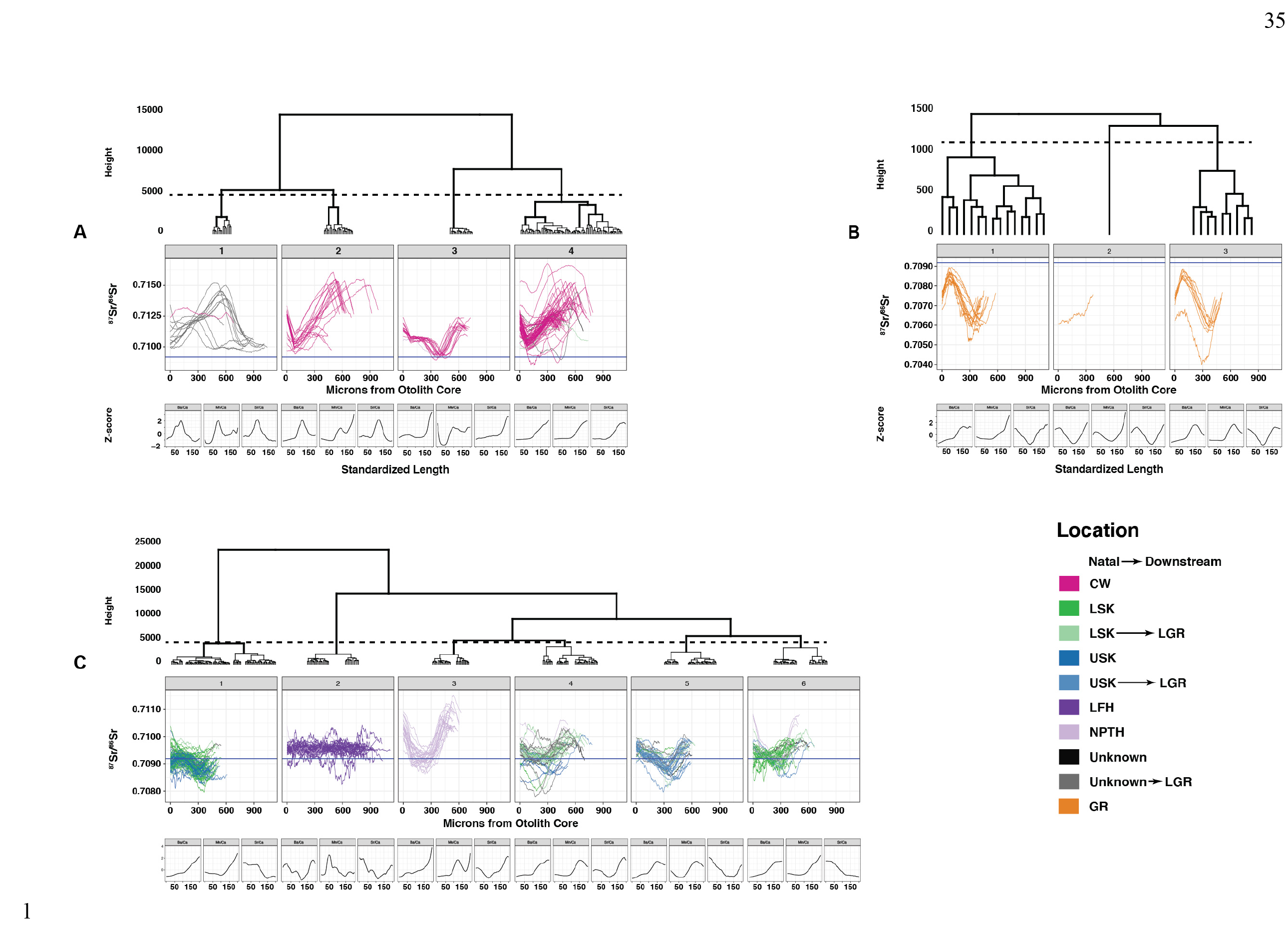
Multivariate, hierarchical DTW sub-clustering of juvenile fish using ^87^Sr/^86^Sr, Sr/Ca, Ba/Ca, and Mn/Ca transects are shown. Panels A through C represent the sub-clustering for each cluster based on overall transect mean. Dashed lines show the location the dendrogram was cut to determine the cluster solution. Transects are colored by the known location of the fish. Some fish were captured in their natal location, released, and recaptured downstream in LGR. Small panels below each cluster show the z-normalized centroid for each trace-element in each cluster. The blue horizontal line represents the global marine value of ^87^Sr/^86^Sr (0.70918) for reference.

**Table 3.**
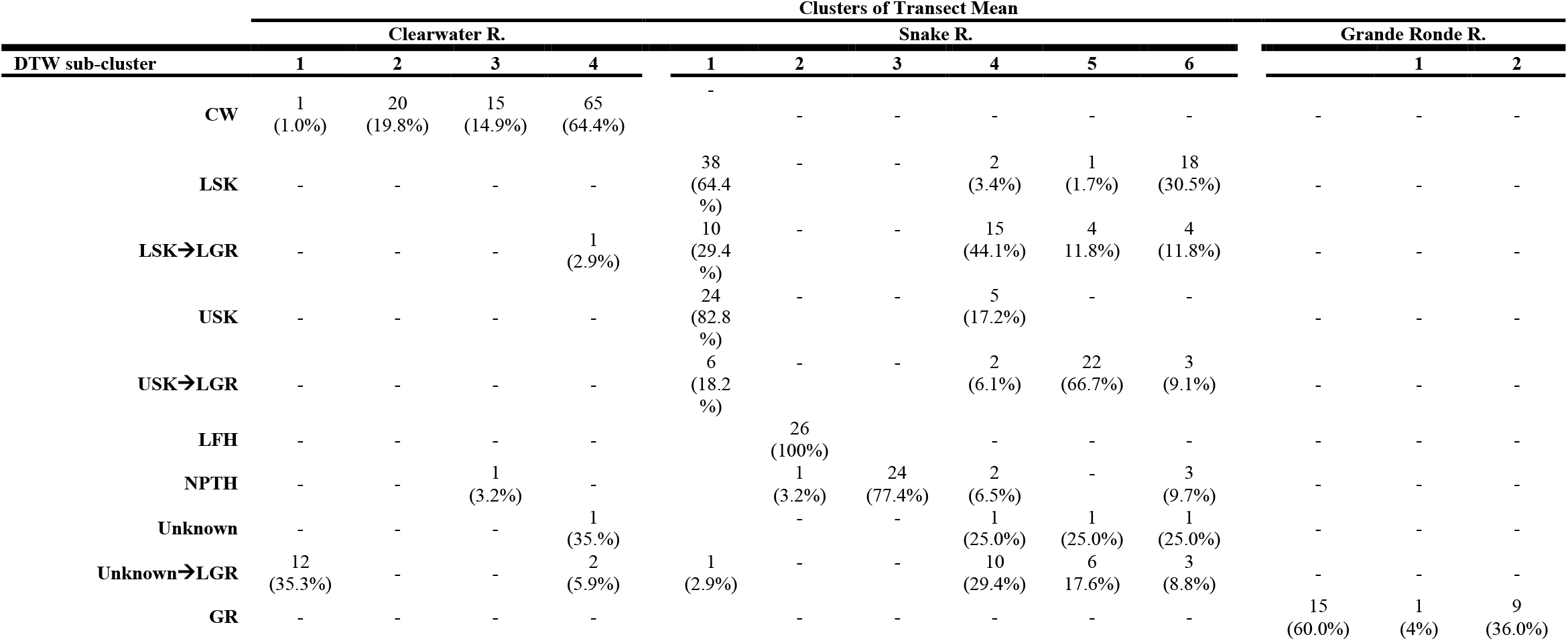
Results of two-level clustering of known-origin juvenile fish using multivariate DTW distance. Fish were first clustered by the mean 87Sr/86Sr of the entire transect using k-means, resulting in three broad clusters corresponding to the river of origin (Clearwater River, Snake River, and Grande Ronde River). These clusters were sub-clustered using hierarchical clustering and multivariate dynamic time warping distance of 87Sr/86Sr, Sr/Ca, Ba/Ca, and Mn/Ca signatures on the transect across each otolith. Sample size in each group is shown, with the percentage of each known-origin group shown in parentheses.

Sub-cluster 4 in the Clearwater River cluster contained fish predominantly of CW origin, with 2 fish of unknown natal origin, one fish of unknown origin captured in LGR, and 1 with LSK natal origin captured in LGR. The ^87^Sr/^86^Sr transects for Cluster 4 showed a similar sharply ascending pattern as Cluster 2. However, they were distinguished by an ascending pattern in Ba/Ca, Mn/Ca, and Sr/Ca, while Cluster 2 displayed a sharp peak in Ba/Ca and Sr/Ca. The reasons for the differences in the shape of the elemental ratio transects are unknown, but it is possible that this represents a meaningful difference in life-history despite the similarity in ^87^Sr/^86^Sr profile.

Sub-clustering of the Grande Ronde cluster using DTW resulted in three clusters. Clustering height was an order of magnitude lower than for the other groups, indicating that clusters were more closely related (Figure 5B). Sub-clusters 1 and 3 displayed very similar ^87^Sr/^86^Sr transects but were distinguished by increasing Ba/Ca and Mn/Ca in sub-cluster 1 and a peak in Ba/Ca and Mn/Ca midway through the transect for Cluster 3. Cluster 2 was made up of a single fish which displayed a unique upward trending ^87^Sr/^86^Sr signature but similar elemental ratio patterns to sub-cluster 3. All of the fish in the Grande Ronde clusters were captured in the GR (Table 3).

Sub-clustering of the samples from the Snake River cluster using multivariate DTW resulted in 6 well-defined clusters. Cluster 1 displayed a descending pattern of ^87^Sr/^86^Sr ratios beginning near the global marine signature and declining toward the ^87^Sr/^86^Sr signature of the Upper Snake river before rising slightly toward the global marine signature near the end of the transect (Figure 5C). Elemental ratios showed an increasing trend in Ba/Ca and Mg/Ca, with a decreasing trend in Sr/Ca. The cluster contained a majority of fish with natal origins in LSK, with 38 captured in LSK and an additional 10 initially captured in LSK before being recaptured downstream in LGR. Fish from the USK made up the remainder of the known origin fish in this sub-cluster, with 24 captured in the USK and 6 initially captured in the USK before being recaptured in LGR.

Snake River sub clusters 2 and 3 were comprised almost entirely of fish captured at each of the two hatcheries in the study (Figure 5C). Snake River sub-cluster 2 showed a largely flat ^87^Sr/^86^Sr transect and highly variable elemental ratios. It was made up of 26 fish from LFH, comprising all of the fish from this hatchery included in the study. A single fish from NPTH was also included in this cluster. Sub-cluster 3 was similarly made up of hatchery origin fish, with 24 fish from NPTH comprising the only members of the group.

Snake River sub-clusters 4 and 5 were comprised mostly of downstream migrants captured in the LSK and USK respectively, later re-captured at LGR (Figure 5C, Table 3). Sub-cluster 4 was dominated by fish originating in LSK before being recaptured at LGR (15). The ^87^Sr/^86^Sr transects followed a pattern originating near the global marine signature and increasing toward the signature of the LSK river. Ba/Ca transects showed an increasing pattern, Mn/Ca exhibited a peak at 150 cells in the re-interpolated data, and Sr/Ca showed an increasing pattern with a peak near the end of the transect. In contrast sub-cluster 5 was made up largely of fish originating in USK before being recaptured in LGR (22). This group displayed a pattern of ^87^Sr/^86^Sr ratios decreasing below the global marine signature, toward the signature of the USK, before rising toward the LSK signature at the end of the transect. Elemental ratios in this group showed similar patterns to sub-cluster 4 in Mn/Ca but decreasing Sr/Ca and a late peak in Ba/Ca. Snake River sub-cluster 6 contained a majority of fish captured in LSK (18), with fish from LSK captured in LGR (4), USK captured in LGR (3), and NPTH (3) (Table 3). This group displayed an ^87^Sr/^86^Sr transect increasing from the global marine signature toward the LSK signature, with Ba/Ca and Mn/Ca increasing across the transect and Sr/Ca decreasing (Figure 5C).

Overall, multivariate DTW provided a clustering solution that clearly identified known life histories, while also identifying additional life-histories within the data which other methods did not. Within the Clearwater River DTW sub-cluster 1 contains fish captured at LGR with unknown natal-origin. The change from an increasing ^87^Sr/^86^Sr pattern consistent with the CW early in life, to a lower LGR signature later in life is evident (Figure 5A). These unknown-origin fish were collected by USGS in 2012 in the forbay of Lower Granite dam and the collection notes include, “Unknown origin. Likely from Clearwater but could be hatchery or natural,” based on the expected timing of outmigration from the Clearwater River. This provides evidence that the DTW algorithm is able to successfully match these fish to their natal location, despite the change in transect shape caused by movement into the lower ^87^Sr/^86^Sr signature of LGR. Further, three of these fish demonstrate movement into LGR at a much earlier point than the others (Figure 5A), yet the time-warping nature of DTW and the similarity of their elemental ratios allows them to be clustered together.

Snake River sub-clusters 1, 4, 5 and 6 support the finding from the multivariate DFA that USK and LSK fish are confounded due to early movement (Figure 5C). Each of these clusters ^87^Sr/^86^Sr transects originate near the global marine signature in the maternally influenced region of the otolith (∼0-150μm, Hegg et al. 2018). Transects them toward either the signature of the USK which is below the global marine signature (sub-cluster 1 and 5) or the USK and LGR which is largely above the global marine signature (clusters 4 and 6) during the natal period (∼250-400μm, Hegg et al. 2018). The shape of the ^87^Sr/^86^Sr transect due to movement downstream to LGR is clearly visible in cluster 5, and the addition of elemental transects clearly delineates this cluster from a similar ^87^Sr/^86^Sr shape for fish originating in the LSK (cluster 4 and 6). Sub-cluster 1, however, contains a majority of fish whose transects clearly end within a signature lower than the global marine signature which should indicate an origin in the USK. Despite this the majority of fish in this sub-cluster were captured in LSK. This indicates that these fish likely originated in the USK, moved downstream to the LSK, and were captured before their signatures had equilibrated to the LSK signature.

### Combining DTW with Discriminate Function Analysis

Multivariate hierarchical DTW clustering of the confounded USK/LSK samples in the discriminate function training-set was done to test the ability of DTW to separate the confounded group. This clustering of the confounded USK/LSK group resulted in three clear clusters (Figure 6A and B). The first cluster contained fish originating in LSK (83%) with a shape similar to cluster 6 in the multivariate sub-clustering above. The second cluster contained a mixture of fish captured in the LSK (51%) and USK (45%), but with an ^87^Sr/^86^Sr shape similar to Snake River multivariate sub-cluster 1 above. The third cluster contained downstream migrants recaptured in LGR after being initially captured in the USK (44%) and LSK (41%) before being recaptured in LGR. The dendrogram did not provide simple separation of this downstream migrant group into LSK-LGR and USK-LGR clusters. This result indicated the ability of DTW to separate LSK fish from USK fish, as well as fish from each natal region captured downstream at LGR, based on the different transect shape produced in each natal river. Sakoe-Chiba window width did not have a large effect on clustering results.

**Figure 6-.**
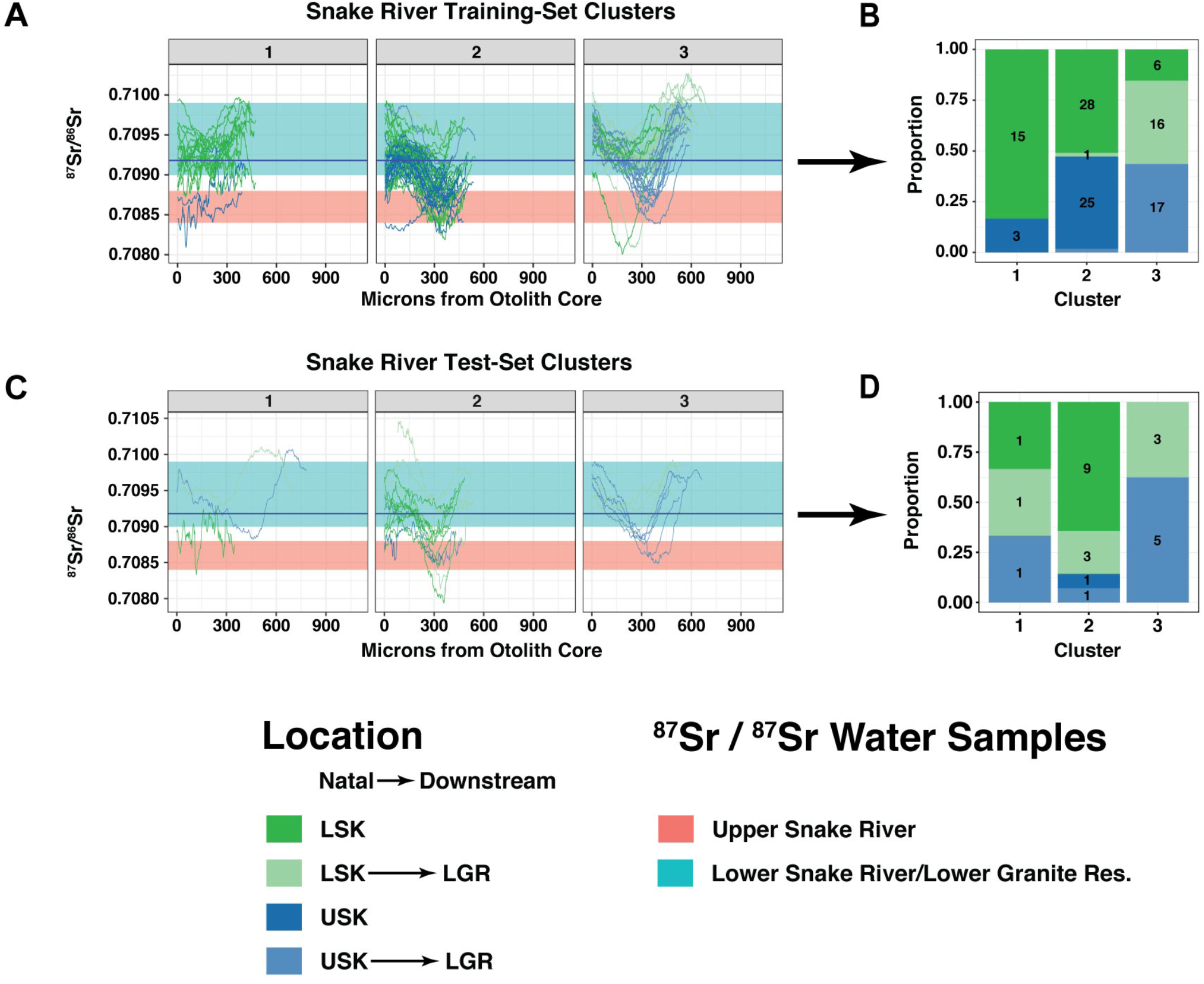
Multivariate, hierarchical DTW clustering of juvenile fish in the confounded USK/LSK group from the discriminate function are shown. Clustering was performed on ^87^Sr/^86^Sr, Sr/Ca, Ba/Ca, and Mn/Ca transects. Panel A shows the clustering of fish in the discriminate function training set. The proportion of fish in each training-set cluster is shown in Panel B, with the number of fish in each cluster shown within the colored region of the plot. Panel C shows the clustering of fish in the test set. The proportion of fish in each test-set cluster is shown in Panel B, with the number of fish in each cluster shown within the colored region of the plot. Fish transects are colored by their location of capture. Some fish were captured, released, and subsequently re-captured downstream in the LGR. Colored rectangles show the range of water samples collected over a longterm study of the Snake River spawning areas (Hegg et al. 2013a, 2018). The blue horizontal line represents the global marine value of ^87^Sr/^86^Sr (0.70918) for reference.

Testing this classification using 1-nearest neighbor to assign group membership resulted in the majority of the test set being assigned to the second cluster (Figure 6C and D). Cluster one contained three fish, one captured in LSK and two downstream migrants captured in LSK and USK respectively and recaptured in LGR. Cluster 2 contained nine fish captured in LSK, three downstream migrants originating in LSK and recaptured in LGR, one fish captured in USK, and one downstream migrant from USK captured in LGR. Cluster 3 was made up entirely of downstream migrants with three fish originating in LSK and five originating in USK before being captured in LGR.

Comparison of the cluster transects to the range of water ^87^Sr/^86^Sr signatures collected from the USK and LSK reaches supports the contention that clusters 1 and 2 separate fish by their true natal river reach (Figure 6). Cluster 1 mostly contains fish with a ^87^Sr/^86^Sr signature reflective of the LSK in the natal region between ∼250µm - 450µm from the otolith core in both the training and test set, and most were captured in the LSK reach (Figure 6) Cluster 2 shows a signature below the global marine value, reflective of the USK reach water samples, during the natal period (∼250µm - 450µm). This cluster is split between fish captured in the USK and LSK, despite the samples having a consistent shape. This similarity in transect shape provides additional evidence that early downstream movement is confounding the ^87^Sr/^86^Sr signatures of juveniles captured in the LSK reach. Cluster 3 is made up of a mix of fish from both LSK and USK natal origins which subsequently display an increasing signature consistent with the LSK water samples prior to capture, consistent with their known downstream migration.

The test-set results indicate that the clusters are robust to transect shape, with test-set transects largely mirroring the ^87^Sr/^86^Sr transect shapes from the training set. Few test-set fish were assigned to cluster 1, making evaluation difficult. Cluster 2 appeared to robustly contain fish with a transect shape indicating USK natal origin. Cluster 3 again contained only downstream migrants captured in LGR, indicating that this cluster was robust in identifying the downstream migrant life-history.

## Discussion

Finding analytical methods that best “extract and abstract those fine-scale features that have relevance…in other scales,” is particularly important in ecology (Levin 1992). Our results indicate that DTW clustering, leveraging the structured nature of time series data, can distinguish important groupings in otolith life-history data. The ability of DTW to efficiently use the additional richness in time series datasets, in effect extracting important fine scale features without aggregation data loss, may be useful for a variety of other ecological datasets as well.

DTW was very sensitive to life-history differences recorded in otolith transects. Univariate ^87^Sr/^86^Sr data was capable of classifying fish to location in ways broadly comparable to the traditional discriminant function technique using multivariate chemical signatures. Further, DTW identified additional life-history patterns that were not apparent in the multivariate discriminate function. The classification of two clearly different life-histories within the Clearwater River group, including a life-history which were subjectively identified as different and held out of the discriminate functions analysis, was particularly interesting (Cluster 2, Figure 4A). This indicates that the use of otolith transect shape may provide a more robust, data-centered, method to identify outlying groups rather than simply relying on expert opinion.

However, univariate DTW clustering was largely unable to reliably distinguish hatchery fish in the Snake River group, mixing NPTH samples with LSK samples due to a similarity in shape, and clustering LFH fish throughout the remaining clusters (Figure 4C). Further, univariate DTW clustering did not perform differently than clustering based on Euclidian distance, which is somewhat surprising. The outmigration timing of juvenile Fall Chinook varies significantly both year-to-year and between individuals within a natal reach (Connor et al. 2005, 2013, Tiffan and Connor 2012). This variation in timing would be expected to negatively affect the Euclidean distance measure. Standardizing the length of each time series and clustering first on the mean may have improved the similarity in shape, or temporal warping may be low enough to make Euclidean distance a viable measure in this population. However, in more complex matching tasks where much more variation and complexity is the norm, such as otolith data of adult fish or searching for specific movements within a larger otolith transect, it is likely that the temporal flexibility of DTW would be superior.

Multivariate DTW clustering was much more successful and sensitive in classifying fish life-history. In the Snake River group multivariate DTW was able to cluster hatchery fish from NPTH and LFH with a high degree of precision (Clusters 2 and 3, Figure 5C). Further, despite the confounding effect of downstream movement DTW appeared to separate fish by USK and LSK natal origin with a high degree of specificity to transect shape (Clusters 1 and 4, Figure 5C). Further, DTW was able to distinguish downstream movement from each of these natal locations into LGR and cluster them separately based largely on differences on differences in the shape of trace-element transects between the upper river and LGR (Clusters 5 and 6, Figure 5C).

The ability of DTW to distinguish more subtle life-history differences is also very interesting. Both in the Clearwater and the Grande Ronde groups DTW identified clusters which appeared subjectively similar based on ^87^Sr/^86^Sr, the primary signature used to infer downstream movement. But underlying differences in the shape of the trace-element transects resulted in these fish being clustered separately. In the case of the Clearwater River group these clusters were relatively far removed on the dendrogram (Clusters 2 and 4, Figure 5A), while in the Grande Ronde group they were more closely related (Clusters 1 and 3, Figure 5B).

Trace-elements in the otolith vary not only in response to differences in concentration in the surrounding water, but also in response to differences in temperature, growth rate, and other metabolic processes (Campana 1999, Walther and Limburg 2012, Limburg et al. 2018). This indicates that while the ^87^Sr/^86^Sr transect may not show differences in life-history, the transect shape of other trace elements may point toward important differences in the life-history or spatio-temporal interaction with the surrounding environment in some sub-groups of fish with otherwise similar downstream movement patterns.

The results of model-based discriminate function analysis show that the traditional approach, aggregating data into a mean from the natal period on the otolith, is effective (Table 1). However, the inability of the multivariate discriminate function to discriminate juveniles from the USK and LSK reaches of the Snake River provides an interesting example of the loss of information due to aggregation.

The water signatures between the USK and LSK reaches are significantly different over multiple years of sampling (Hegg et al. 2013a, 2018). In this case, early moving fish may be identified as LSK when, in fact, they are recent migrants from the USK reach whose chemistry has not equilibrated, or whose natal period does not match the expected 250-450μm location on the otolith due to variation in growth rate. Aggregating this data into a mean value incorporates these erroneous signals and ignores information contained in the temporal structure of the otolith data. Multivariate DTW is able to take the temporal dimension of this data into account, in a quantitative way, to separate the confounded LSK/USK groups (Figure 5C). The fact that natal signatures within each DTW group match the expected water chemistry of the USK and LSK reaches (Figure 6) provides strong evidence of the usefulness of the DTW method. These results indicate that DTW incorporates the fish movement, growth, and chemical data in the otolith to uncover meaningful associations in the data that traditional methods cannot.

The advantages of DTW as an analytical tool might be most apparent when combined with existing analysis techniques. For the data presented here, discriminant function analysis excels at pinpointing fish location based on the highly accurate chemical means that are aggregated from the natal signature. However, in the case of the confounded USK/LSK samples, using the temporal structure of the otolith data through DTW allows this problem to be resolved (Figure 6).

The possibilities of DTW extend beyond the methods presented here. While our analysis is limited to clustering short time series with equal lengths, DTW is capable of more flexible pattern matching. For example, by relaxing the constraint that the time series be of the same length it is possible to search for short, prototype time series within longer series (Tormene et al. 2009, Rakthanmanon et al. 2012). This could be used to find specific short-term fish behaviors, perhaps transitions between specific rivers, within the longer transects of adult fish. In other contexts, this could identify specific hydrological events within many years of hydrograph data, or specific patterns of phenology across years or across a landscape. This “open-ended” method has been used successfully in several examples of extremely large datasets (Tormene et al. 2009, Rakthanmanon et al. 2012). The method is also easily applied to data that is not strictly temporal but can be sequentially ordered, for species identification by shape or identification of movement patterns (Ueno et al. 2006, Cope and Remagnino 2012, Jouary et al. 2016). Also, where good training data exist DTW combined with nearest-neighbor classification has been shown to be a robust and accurate classification method (Kate 2015). Further, DTW can be applied to multivariate time series, though careful pre-processing is required (Mueen and Keogh 2016).

Despite the advances in DTW methods, and demonstrated utility in other fields, ecology has not embraced the technique. These methods are increasingly easy to utilize, with DTW packages available in multiple popular platforms including R, Python, Java and SAS (Leonard and Wolfe 2001, Salvador and Chan 2007, Albanese and Visintainer 2012, Gulzar 2015). Analysis of ecological data is always a balance between detail and parsimony, and often temporal in nature. DTW provides an additional tool for ecologists to maximize the information available to answer ecological questions by taking advantage of the information contained in sequentially structured data.

## Acknowledgements

Thank you to the Kennedy LIFE lab; K. Gillies-Rector, A. Anderson, and N. Wingerter for their insights into the early drafts. In particular, E. Benson and J. Reader deserve credit for prepping a large number of juvenile otoliths for this project. Thanks to the lab undergraduates; A. Miera, J. Greenwood, K. Wilcox, and B. Carman for their help with data post-processing. Thanks also to L. Rayala for early editing.

