## Appendix A for "Let’s Do the Time Warp Again: Non-linear time series matching as a tool for sequentially structured data in ecology"

#### Appendix A – Interpolation of Transects

Interpolation can potentially create artifacts in the data. Similarly, the stretching or compression of the series relative to others could have important effects on the DTW results. Here we show a plot of fifteen randomly selected transects before interpolation, and the same transects after interpolation to 200 cells (Figure 1A, next page).

Most transects look very similar after interpolation. The exception is one fish from the CW (red line), and one from NPTH, which lose the downward trend at the end of the otolith. This short downward trend is likely due to the fish moving into a lower section of the river, a section with a lower ^87^Sr/^86^Sr signature, immediately before capture. Fish with this pattern clustered together with fish who had completed this transition to lower ^87^Sr/^86^Sr, indicating that this loss of the final point in the series did not adversely affect matching.

(figure next page)

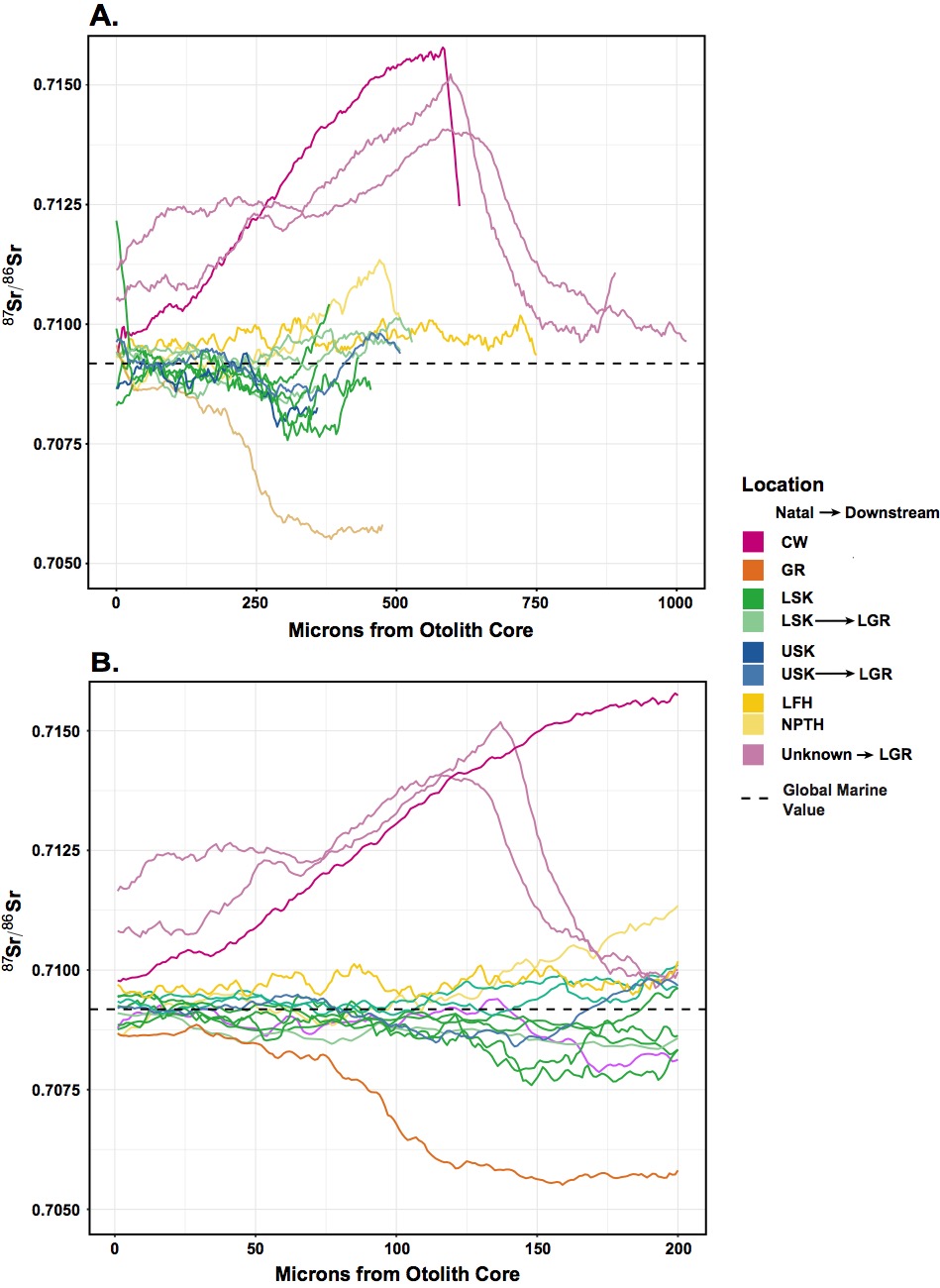

**Figure 1A – Transects Prior to Interpolation**

Fifteen randomly selected transects (**A**) from the study were selected to demonstrate the effect of interpolation (**B**) on the quality of the data. Transects are colored by the known location of the fish. The horizontal dashed line represents the global marine value of ^87^Sr/^86^Sr (0.70918).
